# A “Tug of War” maintains a dynamic protein-membrane complex as shown in all-atom simulations of C-Raf RBD-CRD bound to K-Ras4B at an anionic membrane

**DOI:** 10.1101/181347

**Authors:** Zhen-Lu Li, Priyanka Prakash, Matthias Buck

## Abstract

Association of Raf kinase with activated Ras triggers downstream signaling cascades, towards regulating transcription in the cells’ nucleus. Dysregulation of Ras: Raf signaling stimulates cancers. We investigate the C-Raf RBD and CRD regions when bound to oncogenic K-Ras4B at the membrane. All-atom molecular dynamics simulations suggest that the membrane plays an integral role in regulating the configurational ensemble of the complex. Remarkably, the complex samples a few states dynamically, reflecting a competition between C-Raf CRD and K-Ras4B- membrane interactions. This competition arises because the interaction between the RBD and K-Ras is strong and the linker between the RBD and CRD is short. This study reveals a mechanism that maintains a modest binding for the overall complex at the membrane to facilitate fast signaling processes. It is likely a common mechanism for other multi-protein, if not multidomain proteins at membranes.

## Introduction

The regulation of peripheral membrane protein function is achieved largely by the mutual interactions between the multiple proteins involved, but may be heavily influenced also by the interactions the proteins make with the membrane. While the former can be characterized in part by experiments in solution, resolving the structure of a protein complex at a membrane remains a formidable challenge experimentally, given the nature of interactions involved. Recently techniques such as cryo-electron microscopy, solution NMR with nanodiscs, and EPR/fluorescence correlation methods of proteins bound to liposomes are advancing the characterization of the interfaces, of the orientation and of the dynamics of peripheral membrane proteins at membranes (Calvez et al., 2016; Denisov et al., 2016; Karandur et al., 2017; Mazhab-Jafari et al., 2015; Pérez-Lara et al., 2016). However, computational methods, such as dynamics simulations, are becoming particularly powerful in modeling the structures of proteins at membranes (Gorfe et al., 2007; Grauffel et al., 2013; Ryckbosch et al., 2017; Yamamoto et al., 2016). In this study we apply this latter approach, specifically all-atom molecular dynamics, to our knowledge for the first time to an oligomeric Ras – effector protein complex at a membrane.

Members of the Ras family of small GTPases are anchored to the intracellular leaflet of the plasma membrane and are a key regulator of cellular signal transduction: they convert signaling inputs from multiple transmembrane receptors to downstream activation, typically of kinases, eventually reaching and activating transcription factors in the cells’ nucleus (Simanshu et al., 2017). Signal transmission is achieved by the activation of Ras, converting it from Ras.GDP (inactive) to Ras.GTP (active) with the help of guanine nucleotide exchange factors, or GEF proteins. Activated Ras triggers down-stream signaling through several pathways, including the Raf-MEK-ERK cascade. Oncogenic mutations, which usually result in permanently activated Ras and its binding to Raf, lead to severe cellular dysfunction. Importantly, ~20-30% of all human cancers harbor an oncogenic Ras mutation (Prior et al., 2012; Schubbert et al., 2007). Strategies that aim to disrupt the several steps which are required for Ras activation, most notably drugs that interrupt the Raf-Ras interaction, are being developed in the recent five years and are expected to be promising therapies in cancer treatment (e.g. Athuluri-Divakar et al., 2016; Ostrem et al., 2013; Welsch et al., 2017).

Downstream effectors such as Raf and PI3K interact with Ras (Mott et al., 2015). In the present investigation we studied the conformational and orientational dynamics of the C-Raf^RBD-CRD^: K-Ras complex bound to a membrane model. That the <u>R</u>as <u>B</u>inding <u>D</u>omain, or RBD (res. 56-132), makes direct contacts with the membrane-bound Ras is well established and structures for binding of the K-Ras4B homologous H-Ras GTPase to Raf have been presented (Aramini et al., 2015; Fetics et al., 2015). There is as yet no study that clearly shows a direct interaction between Ras and the <u>C</u>ysteine <u>R</u>ich <u>D</u>omain, CRD (res. 137-187) which follows the RBD in the sequence Raf, although a number of studies have indicated such interactions (Brtva et al., 1995; Clark et al., 1996). In addition, the CRD was also identified as a membrane binding protein, especially for the lipid headgroup of phosphatidic acid (PA) and phosphatidylserine (PS) (Hekman et al., 2002; Improta-Brears et al., 1999; Mott et al., 1996). It has long been known that in most cases membrane localization is required for Ras activity. Specifically, the membrane helps to locally concentrate Ras proteins and likely directs Ras oligomerization, Ras cluster formation as well as association with other proteins, including Raf (Abankwa et al., 2010; Weise et al., 2011; Zhou et al., 2017). Recent experiments as well as computer simulations have shown that the cell membrane determines the orientational preference of Ras relative to the membrane (Kapoor et al., 2012; Li et al., 2017; Mazhab-Jafari et al., 2015; Prakash et al., 2016a), which is predicted to have an effect on Raf: Ras recognition events. Despite the many studies on the interactions of isolated Raf or Ras domains with the membrane, a biophysical-structural study of Raf: Ras as a protein complex at the membrane has not yet been reported, neither with experimental nor computational methods.

How are Raf-membrane, Ras-membrane and Raf-Ras interactions integrated together in order to determine the structural features and function of the Raf: Ras complex at the intracellular membrane leaflet? In the present research, computational studies provide a powerful avenue for the detailed structural and dynamic studies of the C-Raf^RBD-CRD^: K-Ras4B complex at the membrane. The observations lead to the formation of further hypotheses, suggest key residues for mutagenesis for future functional studies and provide a mechanism of that can generalized to other protein complexes at membranes.

## Results

### Configurations of C-Raf

We performed five independent all-atom MD simulations of the GTPase membrane anchored complex of C-Raf^RBD-CRD^: K-Ras4B for 1 μs each (Fig. 1). Before this, we first examined the configurations of an isolated C-Raf^RBD-CRD^ (denoted C-Raf from now) in solution with three independent simulations of 1, 0.5, 0.5 μs respectively (Fig. 2a). The configurations sampled a few major clusters (Fig. 2b-c; Fig. S1a). Overall, the CRD samples a wide configurational space relative to RBD (the figure shows a superposition on the RBD), suggesting a large flexibility of the linker between the two domains. But it is noticeable that the CRD cannot reach the β-sheet surface (β2, β1 and β5) and the region (α1 and β2) of the RBD that is used for Ras binding. Therefore, none of these C-Raf configurations are likely to have major clashes with the binding of K-Ras. The program FPModeller (Pham et al., 2007) independently predicts a similarly wide configurational space of C-Raf and similar excluded regions, by rotating residues in the short linker (132 to 137) (Fig. S1b). The interdomain linker is short and this makes the CRD less easy to access the distal α1-β2 region of the RBD. However, there is also a mismatch in surface electrostatic potential between these regions (of the α1-β2 region as well as the β2-β1-β5 region) with the CRD surface, making such configurations unfavorable (Fig. 2c, lower). In contrast, interactions between the CRD and the RBD loops between β4 and α2, β3 and α1 as well as between β1 and β2 of the RBD are occasionally established (Fig. S1c), which contribute to the higher population of configurations Cluster #1, Cluster #2 and Cluster #3 (Fig. 2b).

**Figure 1:**
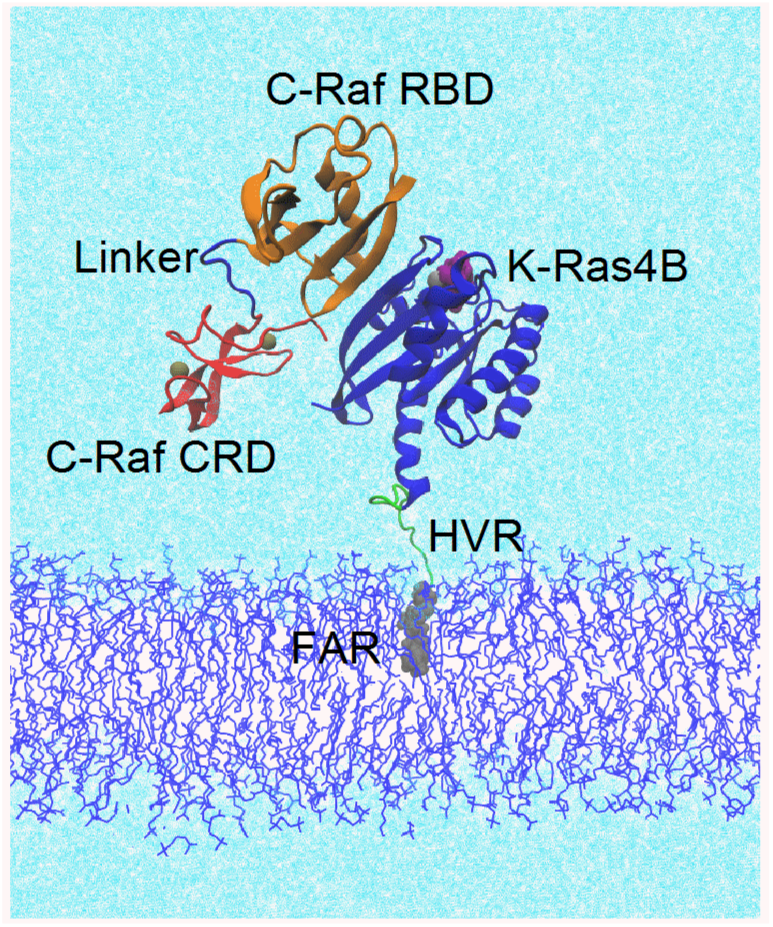
C-Raf^RBD-CRD^: K-Ras4B complex at a mixed membrane consisting of 80% POPC and 20% POPS. The K-Ras4B protein consistes of the globular, catalytic domain (CD, res. 1-166) and the largly unstructured hypervariable region, HVR (res. 167 to 185). C-Raf comprises the Ras Binding Domain, RBD and Cysteine Rich Domain, CRD, connected by a short linker. Proteins shown as mainchain cartoon; K-Ras (blue); RBD (orange); CRD (red); small molecules/ions as space filling: farnesyl group (grey); GTP (purple); Zn and Mg (tan); show as lines: linker region (blue); HVR (green), membrane (blue); water (light cyan).

**Figure 2:**
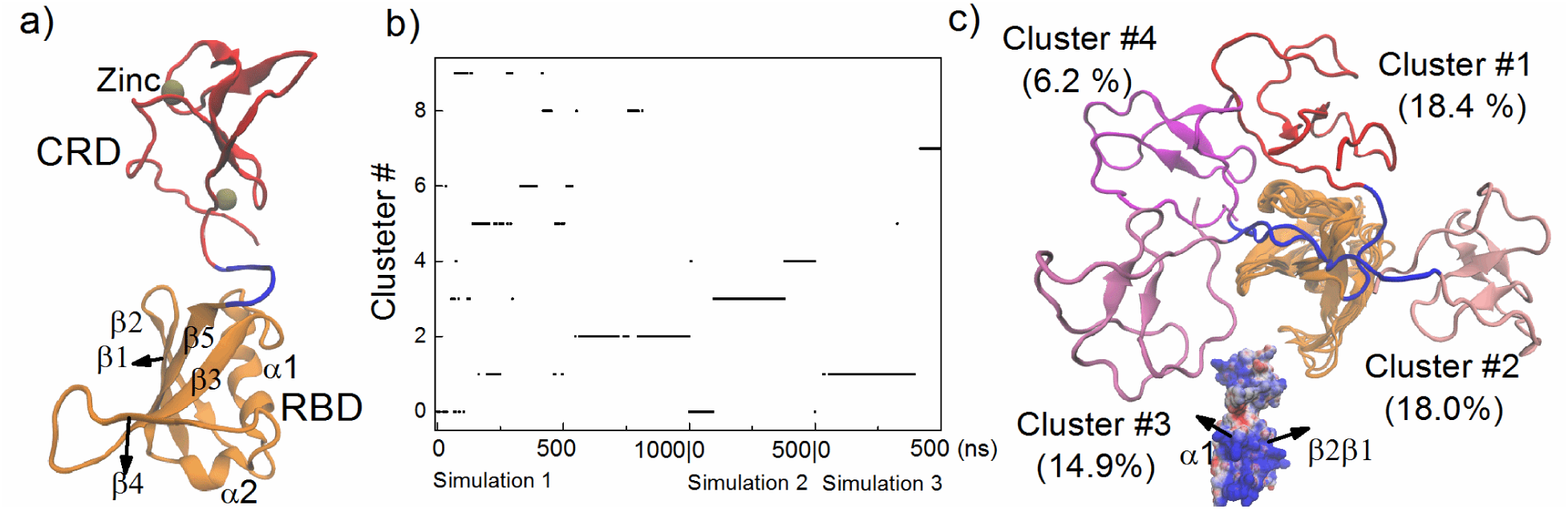
Configurations of C-Raf^RBD-CRD^ (C-Raf) in solution. (a) Starting configuration of C-Raf, (b) Clustering of configurations with an RMSD cut-off distance of 5 Å, (c) Representative configurations superimposed on RBD (pointing away from observer). Cluster #1-4 are shown (others in Fig. S1a). Surface electrostatic potential of domains (seen as in a), rotated by +90 ° around z).

When bound to membrane anchored K-Ras4B, the configurational flexibility of C-Raf is largely preserved (Fig. S2a-b). The configurations are comparable to the configurations of an isolated C-Raf, although the populations of the different configurations are changed to different extents (Fig. S2c). In addition, the radius of gyration of the two domains of C-Raf is more extended when bound to membrane anchored K-Ras4B (Fig. S2d). These differences reflect the additional interactions of C-Raf with K-Ras4B as well as those of C-Raf CRD with the membrane.

### Interaction of C-Raf with K-Ras4B

The C-Raf: K-Ras4B interactions are mostly confined to those that are known from the C-Raf^RBD^: H-Ras crystal structure (Fetics et al., 2015), and this interface is highly persistent in the simulations. The β-sheet interface is established between residue 37-39 of K-Ras4B and residues 65-70 of the RBD (Fig. S3a). Outside these regions contacts K84:E31/D33, V88:I21/Y40 and R89:D38/S39/Y40 are seen between the RBD and K-Ras (Fig. S3a). Overall, these detailed protein-protein contacts are in close agreement with the interactions seen in the experimental structure (Fetics et al., 2015). Two reports also suggested a direct interaction between the CRD and the switch II region of Ras. GTP (Aramini et al., 2015; Brtva et al., 1995). These interactions are not observed, however, in any of the 5 simulations (Fig. S3b). We performed one additional simulation (#6) by placing the CRD at the switch II region. In fact, over the course of the simulation the CRD gradually moves away from the switch II region of K-Ras4B (Fig. S3c). Experimentally, the binding of the RBD or of the CRD to Ras were measured separately in the literature. When RBD and CRD are linked, we suggest that the CRD has a low potency for Ras binding, as the CRD needs to orientate itself to an unfavorable position in order to display the α1-β2 region of the RBD for interaction with switch II. In a competition the RBD has a much greater advantage compared to the CRD for binding to Ras given a K_d_ of 20 nM for the RBD versus an approximately 5.5 weaker affinity for CRD (measured for H-Ras in Ghosh et al., 1994). Essentially, the CRD is engaged in a variety of competitive interactions amongst CRD-RBD, CRD-Ras and CRD-membrane contacts, with the latter probably being the most favorable.

### The K-Ras4B catalytic domain and the C-Raf CRD interact with the membrane in a dynamic manner

Both K-Ras4B catalytic domain and C-Raf CRD region are able to contact the membrane in the simulations, but, when bound together as a protein-protein complex, do so in a dynamic way. For convenience of analysis, the K-Ras4B catalytic domain (CD) is further divided into two parts, i. e, CD1 or lobe1 containing residues 1 to 86 (containing P-loop, switch 1, switch 2, and the Raf RBD-binding interface of Ras), and CD2/lobe2 with residues 87 to 166 (containing helix 3, 4 and 5) (Gorfe et al., 2008). Fig. 3 depicts the time evolution of the distance of the K-Ras4B CD2 or C-Raf CRD to the membrane center for simulation #1 and #3 (see Fig. S4 for others). In simulation #1 (Fig. 3a and Fig. 3c), the K-Ras4B catalytic domain binds to the membrane for the first time during the early 100 ns, but later it undergoes a few dissociation-association events. In comparison, the CRD gradually and spontaneously moves toward the membrane in the first 400 ns, and remains bound to the membrane in the next 600 ns. In simulation #3 (Fig. 3b and Fig. 3d), the K-Ras4B catalytic domain reaches the membrane first at ~100 ns. However, along with the movement of the CRD toward to the membrane at ~350 ns, the K-Ras4B catalytic domain then moves away from the membrane. Later, the positions of both K-Ras4B and CRD undergo several fluctuations. Simulations #2, #4 and #5 show a range of similar scenarios (Fig. S4). Overall, except for the CRD in simulation #1, perhaps, the membrane contacts of K-Ras4B or CRD are not highly persistent but are dynamic in nature (see discussion regarding convergence and optimal contacts below). This differs from previously reported situations when isolated K-Ras4B (and also -4A) are placed at a membrane with the same concentration of POPS lipid molecules, where K-Ras is mostly bound to the membrane, detaching less frequently (Li et al., 2017; Prakash et al., 2016a).

**Figure 3:**
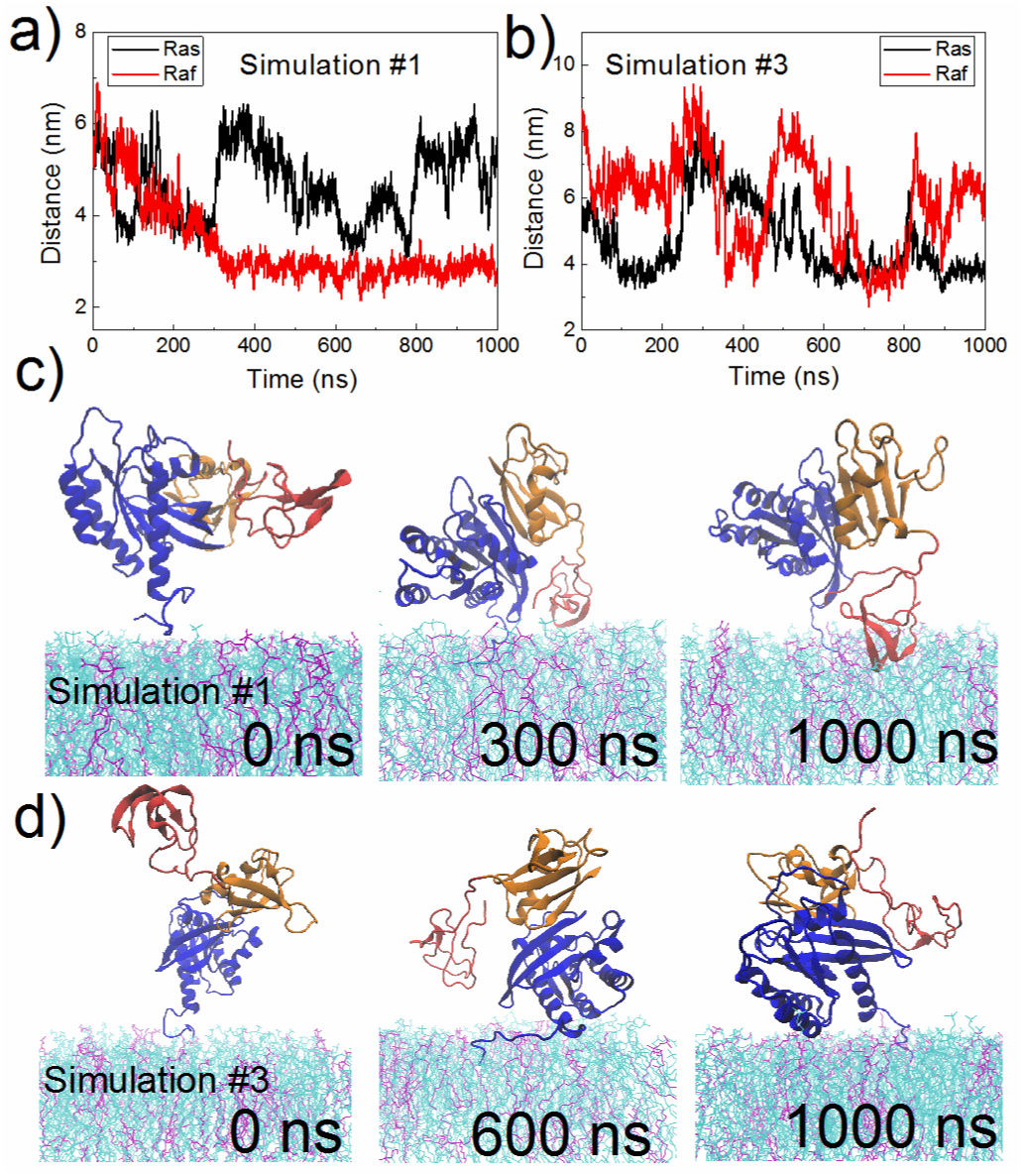
Dynamic membrane binding of C-Raf: K-Ras4B complex. (a-b) Time evolution of the distance of the center of K-Ras4B CD2 (residues 87 to 166) or the C-Raf CRD center to the membrane center. Snapshots taken at various time points from (c) simulation #1 and (d) simulation #3. Color scheme as in Fig.1 except, POPC in cyan, POPS in purple.

### Interface of K-Ras: membrane interaction

Residues involved in contacting the membrane are plotted in Fig. 4a and Fig. 4b for K-Ras4B and C-Raf separately as a function of sequence and as averaged over the 5 simulations. For the catalytic domain of K-Ras4B, the membrane interacting residues belong mostly to helix 3 and helix 4, less so to helix 5 and a small loop segment between β2 and β3 (res. 43-50, loop 3). These interface residues establish the two dominant orientations of K-Ras at the membrane (O3 and O4V; numbering with reference to our previous study; see also Fig. S5). In one orientation, helix 3 and 4 are bound to the membrane; in the other one, the loop 3, and partly helix 5 are the membrane interacting regions. For an isolated K-Ras, the O3 and O1 orientation (where β1-β3 associates with the membrane) are the dominant orientations (Li et al., 2017; Prakash et al., 2016a). When bound to C-Raf, the O1 orientation is completely abolished, as the corresponding β1-β3 interface of K-Ras is now strongly bound to C-Raf. Therefore, O3 becomes the dominant orientation for K-Ras in complex with C-Raf at the membrane. Additionally, the membrane association of the CRD, in some cases, also brings the K-Ras loop 3 close to the membrane, resulting in the popularity of a variant of a previously characterized orientation, O4 (here denoted O4V, where not only the loop 3, but also helix 4 and 5 are largely involved in membrane contacts). As shown in Figure 4c, a modest clustering of anionic POPS lipid molecules is observed under and around the C-Raf: K-Ras4B complex (especially under the GTPase). On average, 18% of the POPS are distributed within 2.5 nm of the center of K-Ras4B CD1 (accounting for ~13.6% membrane area). Such clustering is expected to enhance the binding of K-Ras to the membrane (Li et al., 2017; Prakash et al., 2016a). It should be noted that 1 μs presents an adequate amount of time for PS lipids to re-organize in the POPC/PS bilayer under these conditions (e.g. Li et al., 2017; Perrin et al., 2015), although the simulations may not be completely converged for the more rapid re-orientational transitions and some hysteresis could be present.

**Figure 4:**
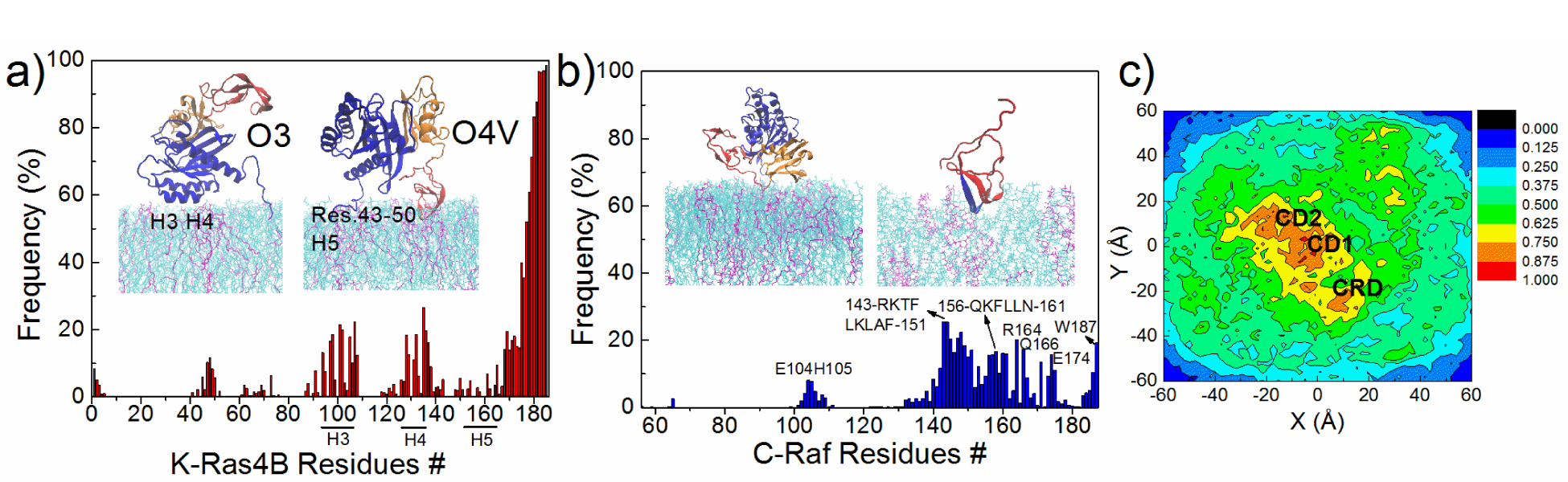
Interface of the K-Ras4B: membrane interaction. (a) Frequency of K-Ras4B - membrane contacts (residues within 5 Å of membrane surface); position of helices 3-5 is indicated. Inset images show two major orientations of complex relative to the membrane. (b) POPS distribution at the membrane. The distribution is normalized to make the maximum 100%. K-Ras4B CD1 is centered at (0, 0), the horizontal displacement between the center of mass of K-Ras4B CD1 and that of C-Raf RBD is aligned to the X axis. The last 500 ns trajectories were used. (c) Frequency of C-Raf - membrane contacts. The region, res. 143-150, is indicated in blue. Inset images: Representative orientations of RBD and CRD at the membrane.

### Interfaces for the C-Raf: membrane interaction

As for C-Raf, only a few residues of the RBD (res. 56 to 132) can contact the membrane (Fig. 4b), when bound to K-Ras. But the RBD-membrane contact frequency is rather low with the largest occupancy about 8.1% and 7.7% for E104 and H105, respectively. The most prominent membrane contacts involve residues in the CRD. Almost all of the residues in the CRD (res. 137 to 187) are able to contact the membrane more or less equally, except for a few residues hidden inside the folded conformation such as residues around V180 (Fig. 4b). The CRD-membrane interaction is driven by both hydrophobic and by electrostatic contacts. The CRD can become partially buried into the membrane (most notably in simulation #1), with hydrophobic residues L147, L149, L159 and L160 largely buried into the membrane. The same region is also predicted as membrane inserted by the OPM (Orientations of Proteins in Membranes) webserver (Lomize et al.; 2006). Positively charged residues including R143, K144, K148, K157 and R164 also interact with the lipid bilayer, which is consistent with the experiment-based report that the 143-RKTFLKLA-150 segment of the CRD has a role in membrane binding (Improta-Brears et al., 1999). In addition, cation-π interactions established between F146, F158 and especially F151 and W187 and the Nitrogen of the lipid headgroup (Fig. S6a), as well as hydrogen bonding interactions between residues T145, Q156 and N161 (Fig. S6b) also aid the membrane adhesion of the CRD.

### Configurations of the C-Raf: K-Ras4B complex at the membrane

The configurations of the C-Raf: K-Ras4B complex with respect to the lipid bilayer are further characterized using several geometric parameters as shown in Fig. 5. In Fig. 5a, the distance of the K-Ras4B catalytic subdomain (CD2) to the membrane center is plotted versus the distance of the CRD to the membrane center, mapped over all 5 simulations. In one dominant conformation the CRD is away from the membrane and the K-Ras4B CD2 is bound to the membrane (D2 ~6-8 nm and D1 ~3.5 nm, respectively). In contrast, another dominant conformation corresponds to CRD approaching the membrane K-Ras CD2 pointing away from it (D2 ~3-3.5 nm and D1 ~4.5-5.5 nm, respectively). Otherwise, there are instances when none of the domains are in contact with the membrane and times when both the K-Ras4B catalytic-domain and C-Raf CRD regions are close to the membrane (~3.5 nm, ~3.7nm, respectively), although such instances are rare. Fig. 5b plots another pair of parameters that characterize the relative position and orientation of the RBD and the K-Ras4B CD1 regions. Again, two orientations dominate: one with the RBD: K-Ras4B CD1 at ~5.3 nm from the membrane center and with the RBD slanting upward (tilt angle centered at ~50°); another represents a position tilted at around 95° and is centered at ~4.2 nm. Unbound states can also be identified where the RBD: K-Ras4B^CD1^ is far away from the membrane and almost being parallel with the membrane surface. Overall, we classify the well populated configurations of the C-Raf: K-Ras4B complex at the membrane into four possible states based on the variables D1 and D2 (Fig. 5a). Representative conformations for these four states are shown in Fig. 5c. The first two states (CRD+/RAS- and CRD-/RAS+) represent the most dominant configurations for the complex. Moreover, there are large overlaps between different simulations (especially simulation #2-#5, Fig. S7). Thus, different states can interconvert to each other, suggesting an inherent dynamics of the protein complex unit with respect to the membrane.

**Figure 5:**
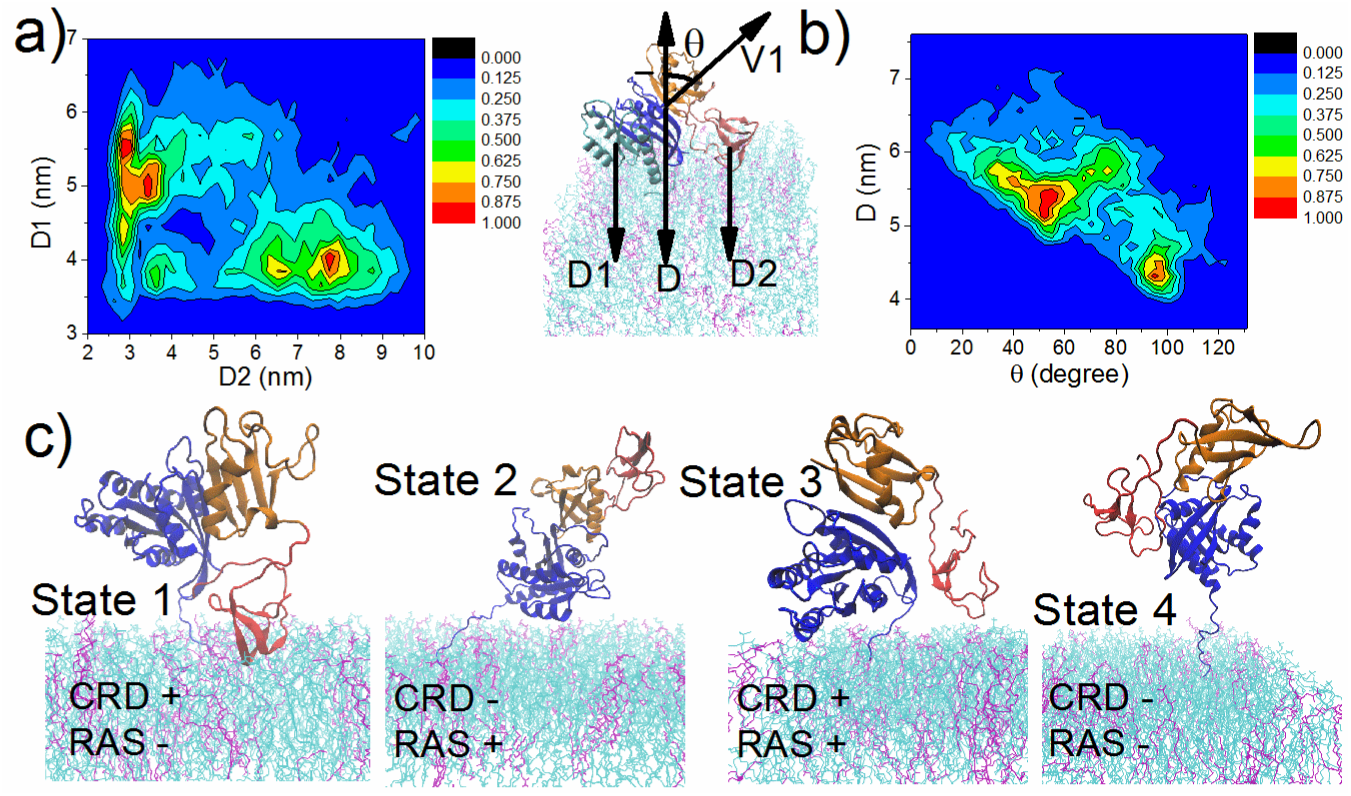
Configurations of the C-Raf: K-Ras4B complex at the membrane. The variables are defined as follows: D1, distance of K-Ras4B CD2, and D2, distance of CRD to membrane center, respectively. D, distance between center of RBD: K-Ras4B^CD1^ and membrane center. V1, vector connecting the center of K-Ras4B CD1 to center of RBD. θ, cross angle between vector V1 and normal to the membrane surface. K-Ras4B CD1, CD2, RBD, CRD in cyan, blue, orange and red respectively. (a) Contour maps (scaled to max.) with variables D1 versus D2. (b) D versus θ. (c) Representative configurations for the 4 possible states of the C-Raf: K-Ras4B complex.

### Incompatibility of CRD (143-RKTFLKLAF-151) and K-Ras4B (helix 3 and 4) in making membrane contacts

It is apparent that the C-Raf CRD and K-Ras4B catalytic-domain contact the membrane mostly in a mutually exclusive manner. The non-cooperative mechanism is largely determined by the topology of C-Raf: K-Ras4B complex. As shown above, the most favorable membrane interaction interfaces are seen as residues 143-RKTFLKLAF-151 for the CRD and helix 3 and 4 for K-Ras4B, respectively (Fig.4a, 4b). However, when helix 3 and helix 4 of K-Ras4B contact the membrane, the RBD as the counterpart of K-Ras4B is moved away from the membrane with a slant angle at ~50° (Fig5b and Fig. 6a). The calculated radius of gyration for K-Ras catalytic domain, RBD, and CRD are 1.5, 1.2 and 1.1 nm respectively (the physical radius is even slightly larger). The distance between center of the K-Ras catalytic domain and the C-Raf RBD is about 2.8 nm and between center of RBD and CRD is averaged at 2.9 nm. Based on the slant angle and the size of these domains, the RBD is at least 3.6 nm away from the membrane surface. Although the CRD has a large freedom to adopt multiple orientations relative to the RBD, none of them enable the CRD to reach the membrane without rotating the K-Ras domain, as the maximal length of the linker plus the CRD is estimated at 2.8 nm (2.9-1.2+1.1 nm, Fig. 6a). Therefore, the most favorable membrane-associated state for CRD (143-RKTFLKLAF-151) and K-Ras4B (helix 3 and 4) cannot coexist. Fig. S8 further discusses the opposite situation when CRD is bound to the membrane, yielding the same outcome. Overall, the C-Raf: K-Ras4B complex may either use 143-RKTFLKLAF-151 of CRD (state 1) or alternatively use helix 3 and helix 4 of K-Ras4B (state 2) to interact with the membrane. In state 3, both K-Ras4B and CRD are close to the membrane, but the complex is not using the 143-RKTFLKLAF-151 region of the CRD and helices 3, 4 of K-Ras4B together in membrane contact, instead each use other, less favorable contacts with the membrane.

**Figure 6:**
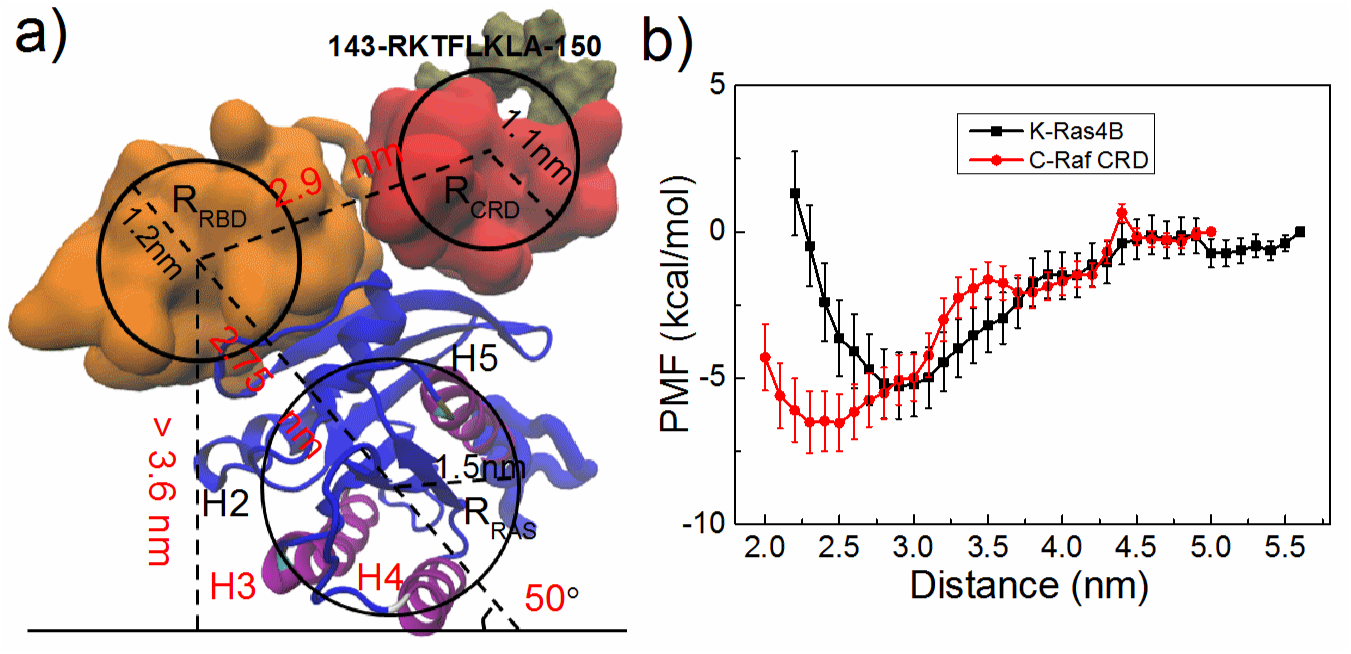
Tug of War between K-Ras4B and C-Raf CRD membrane interactions. (a) Schematic picture of steric/geometric limitations of K-Ras4B and C-Raf complex in different states (here state 2; see SI Fig. S8 for state 1). (b) Potential of Mean Force, PMF, for K-Ras4B and CRD binding to the model membrane. (see also Fig. S10a,b for alignment of sequences for RBD-linker-CRD regions)

### Free energy of K-Ras4B and C-Raf CRD membrane adhesion

We estimate the binding affinity of monomeric C-Raf CRD and monomeric unlipidated K-Ras4B to the model membrane by calculating the <u>P</u>otential of <u>M</u>ean <u>F</u>orce (PMF) along a path of membrane (un-)binding using the most favorable domain orientations (Fig. 6b, see Method). The free energy calculated for monomeric sub-domains are not the exact free energy corresponding to State 1 and State 2 of the protein complex, but their calculation provides a rough estimate for these states. The calculated PMF is -5.3 ± 1.1 kcal/Mol for K-Ras4B binding to the model membrane. The calculated PMF is -6.5 ± 1.0 kcal/Mol for the CRD binding to the membrane. Using the microscale thermophoresis (MST) experiments, for an full-length unlipidated K-Ras4B binding to a membrane comprised of POPC and 5% PIP2 molecules, we measured K_d_ at 23.4 μM (ΔG = -6.31 kcal/Mol) at 298 K (Manuscript in preparation). In the experiment of Gillette et al., a K_d_ was measured at 4.0 μM for lipidated K-Ras4B binding to nanodiscs mixed with DMPC and 30% DMPS. But for unlipidated K-Ras4B, no binding affinity was detected (Gillette et al. 2015). No experimental K_d_ value for CRD binding to the membrane has been reported, but binding to liposomes suggests that the K_d_ should be at least 20 μM (Ghosh et al., 1994). The binding free energy predicted by the OPM sever of -7.31 kcal/Mol, which is larger than the value we estimate. Experimental measurements of the binding affinity of the CRD to the membrane are needed in future. Based on our PMF calculation, the CRD could have a slight but not overwhelming advantage in membrane adhesion compared to the K-Ras4B catalytic domain.

## DISCUSSION

Structural preferences and dynamics of protein complexes at membranes are still a relatively new terrain for discovery. In this study, we chose one complex as an example and performed multiple μs all-atom molecular dynamics simulation for the two-domain fragment C-Raf^RBD-CRD^ (denoted C-Raf) when bound to K-Ras4B at an anionic membrane. While several studies have examined the orientation and dynamics of isolated K-Ras at membranes (Kapoor et al., 2012; Li et al., 2017; Mazhab-Jafari et al., 2015; Prakash et al., 2016a), there are as yet no simulations of C-Raf: K-Ras at membranes. While no experimental structural or biophysical data is available at present for a C-Raf Ras Binding Domain, RBD, Cysteine Rich Domain, CRD, multidomain protein fragment, a relatively tight Ras: Raf RBD association was reported with a K_d_ of 20 nM (Thapar et al., 2004). By contrast the K_d_ for both Ras catalytic domain-membrane and Raf CRD-membrane interactions are estimated at ten to hundreds of μM from our simulations. Thus, membrane binding of the K-Ras4B catalytic domain and of the C-Raf CRD by themselves are only moderately strong, overall consistent with experimental data. One might expect that additional interactions, such as possible direct interactions between the C-Raf CRD and K-Ras4B catalytic domains would substantially increase this affinity via synergistic effects. In fact, this is frequently seen with cell signaling proteins, such as the interactions between WASP, Cdc42 and the membrane, which utilize multiple interactions in a cooperative manner to maximize the signaling output while minimizing output in the absence of coincident input signals (Buck et al., 2014; Dueber et al., 2003). However, this is not the mechanism predicted by calculations here. Importantly, our study predicts a novel competitive mechanism between membrane adhesion of the K-Ras4B catalytic domain and C-Raf CRD of the protein complex (Fig. 5). We are able to rationalize the behavior of this system by considering both the geometric features of the domains within the complex (Fig. 6a) as well as by the estimation of their binding affinity with the membrane (Fig. 6b).

The results from the simulations have several implications for the biological function of the Raf: Ras system: The study confirms that the orientation state (O1), that was previously identified as the dominant state of isolated K-Ras.GTP relative to the membrane (Prakash et al., 2016a; Li et al., 2017), is indeed occluded for Raf-binding. Thus, this state may not be sampled significantly in a cellular environment, given the tight binding between C-Raf RBD and K-Ras (also supposing a high abundance of C-Raf). It is possible that over a long period of time, the structures of K-Ras and the C-Raf RBD may adjust at the membrane to weaken this interaction. However, the fact that the residues involved in this protein-protein interaction as well as the catalytic domain and CRD–membrane contacts are highly conserved (Fig. S9) speaks against this scenario and in any case suggests that the competitive nature of the interactions is likely maintained. Specifically all K-, H and N-Ras are sequence invariant for the first 86 residues, parts of which comprise the contact region with the RBD; on the side of the RBD, res. 65-70 as well as K84, R89 are identical between B- and C-Raf (V88 is changed to M in B-Raf); the region of residues 143-151, primarily responsible for membrane interactions is also identical across many mammalian species (Fig. S9a). The extent of conservation of Ras–membrane interaction sites is also high across the catalytic domains of Ras isoforms (reviewed in Prakash et al., 2016b). Concerning the RBD-linker-CRD region of the protein it is informative to put our simulation results in context of sequence conservation of this Raf protein segment and also with respect to cancer mutations. Fig. S9b shows the alignment of C-Raf with A- and B-Raf homologues. B-Raf is mutated in ~8% of all cancers. ARAF and CRAF mutations are very rare in cancer (Holderfield et al., 2014). The majority of mutations are observed in the activation loop (A-loop) near V600, or in the GSGSFG phosphate binding loop (P-loop) at residues 464–469. Mutation frequency in the linker and RBD-CRD regions is moderate in C- as well as B-Raf.; there are several mutations in the Ras binding interface of the C-Raf RBD and in the regions we find that are interacting with the membrane. However, with exception of L86P and Q156stop, these are all single count according to the COSMIC database. In the linker region, there are two mutations, L136P and R143L with a count of two. The linker region is not changed in length, either between isoforms or between mammalian species. The sequence differs between B-Raf and C-Raf in the linker with the change D132E and H133N (C-Raf numbering); but these changes are not dramatic, so suggesting that the geometric restrictions we observe are likely conserved also between the Raf isoforms, but also between the mammalian species.

The optimal K-Ras4B and C-Raf CRD – membrane bound states present alternative orientations of the overall complex, i.e., membrane contacts of one of them will exclude the best membrane contacts of the other, reflecting a “Tug of War” between membrane interactions of K-Ras4B and C-Raf CRD. This keeps the binding affinity in the modest range of a few μM Kd. Such an affinity is typically associated with cell signaling processes where interactions are kinetically labile and, thus, quick to switch off. An intrinsically modest association gives K-Ras more freedom to interact with multiple regulatory and effectors proteins, which we believe could have a role in the biological function of K-Ras4B. Competition in membrane adhesion may likely be tunable by altering the binding affinity of the individual domains with the membrane. This may be accomplished by posttranslation modifications and/or changes in the membranes’ lipid composition (Zhou and Hancock, 2017). For example, we recently predicted that a PIP2 containing membrane leads to more defined/less dynamic states of K-Ras4A, suggesting that the strength of interaction between the catalytic domain of the GTPase and the membrane are increased (Li et al., 2017).

It is accepted that Ras activity in cells requires its localization to the membrane. Ras clustering is an undisputed observation (Abankwa et al., 2010; Weise et al., 2011; Zhou et al., 2017), but the role of catalytic domain contacts in Ras dimerization in solution and at the membrane is not yet clear. Recently, symmetric Ras: Ras dimers have been predicted through molecular docking, although they mainly represents a weak association (K_d_ ranges from 1 μM to greater than mM) (Muratcioglu et al., 2015; Prakash et al., 2017). Conformational state 2 of C-Raf: K-Ras4B predicted in our simulations does not match any of the predicted dimer forms of Ras (Fig. 5c). But state 1 and of course, the non-membrane binding state 4 are suitable for the formation of Ras dimers, due to the exposure of helix 3 and 4 of K-Ras4B (Fig. 5c) of the dimer interface (Muratcioglu et al., 2015; Prakash et al., 2017). However, in this case optimal K-Ras catalytic domain membrane binding and dimerization would also oppose one another. Nevertheless, CRD-membrane interactions may help to orient the GTPase at the membrane for improved dimerization kinetics, and depending on K-Ras vs. C-Raf concentration, could stimulate K-Ras dimer formation. Similarly, we can only speculate at present, how the two domain C-Raf: K-Ras4B configurations observed in this study may affect the activation of C-Raf Kinase Domain (KD). Higher order C-Raf^RBD-CRD^: K-Ras complexes may have a role in stimulating the dimer formation at the kinase domain. In principle, such complex formation may affect the hinge/loop region that connects the KD and RBD-CRD domains; however this region is rather long (res. 185 to 349) and likely to be flexible, at least in part. Furthermore, the kinase domain of Raf itself is likely to be membrane binding. We are currently working on modeling possible higher order C-Raf: K-Ras complexes, but the added complexity is beyond the scope of this report.

Finally, the findings of our study also have general implications for cell signaling involving protein complexes at the cellular membrane. From other examples it is becoming clear, that the membrane actively participates in the regulation of peripheral membrane protein function (e.g. Calvez et al., 2016; Grauffel et al., 2013; Mazhab-Jafari et al., 2015; Pérez-Lara et al., 2016; Ryckbosch et al., 2017; Yamamoto et al., 2016). However, the protein-membrane interactions typically synergize to compete with protein-protein interactions. For example, the membrane adhesion of Scaffold protein-Ste5, could release an auto-inhibition between its two domains (Zalatan et al., 2012). Another example is the Focal Adhesion Kinase, where both kinase and FERM domains can interact with PIP2 in membranes, leading to a dissociation of these two regions, then allowing the kinase domains to dimerise and activate (Herzog et al., 2017). Now we have found an example of a mechanism involving a membrane peripheral protein complex where domains are in competition by virtue of its interactions with the membrane. The study here reveals this mechanism at molecular level, further adding to the repertoire of signal processes by utilizing peripheral membrane proteins.

## Method

### Method Summary

The model of C-Raf^RBD-CRD^ was built by connecting the crystal structures of the C-Raf RBD (PDB, 4G0N) (Fetics et al., 2015) and C-Raf CRD (PDB, 1FAQ) (Mott et al., 1996) with the native linker. The C-Raf^RBD-CRD^: K-Ras4B complex was further built by docking the modeled C-Raf^RBD-CRD^ structure and to the crystal structure of K-Ras4B (PDB, 4DSO), based on the crystal structure of H-Ras bound with the C-Raf RBD (PDB, 4G0N) (Fetics et al., 2015). The system was placed at a membrane containing 80% POPC and 20% POPS. Five independent simulations were performed each for 1 μs. Umbrella sampling simulations were performed to calculate the free energy of CRD and K-Ras4B binding to the membrane. The CHARMM36m force field was used in all simulations and energy calculations (Huang et al., 2017).

### Starting model: C-Raf^RBD-CRD^ in solution

The structure of the C-Raf^RBD-CRD^ complex was established by assembling the separately available crystal structures of the C-Raf RBD (PDB, 4G0N) and C-Raf CRD (PDB, 1FAQ) using a flexible linker with the protein sequence of residues 133-136 that links these two domains. The C-Raf RBD and C-Raf CRD are connected using the webserver AIDA, where it links the protein subdomains together by addressing the relative orientation between RBD and CRD (Xu et al., 2014). The predicted structure was then used as starting configuration of C-Raf (denoted, SC0). There are two zinc fingers in C-Raf CRD. Each zinc finger has one zinc ion surrounded by three Cysteine residues and one Histidine residue. Histidine is in the uncharged HSE form and Cysteine is patched with CYN (Godwin et al., 2017). The ion coordination within the Zinc finger was stable in all simulations. In total, three independent simulations were performed for 1.0, 0.5 and 0.5 μs respectively.

### C-Raf^RBD-CRD^: K-Ras4B complex at the membrane

The modeled C-Raf^RBD-CRD^ structure (above) was further combined with K-Ras4B (PDB, 4DSO) to construct the initial C-Raf^RBD-CRD^: K-Ras4B complex, guided by a crystal structure of H-Ras bound with RBD (PDB, 4G0N) (Vanommeslaeghe et al., 2012). We carried out 6 independent simulations for the complex, named as Simulation #1 to #6 (see Table S1). In Simulation #1-#4, the center of mass of the catalytic domain of K-Ras4B was placed ~6 nm away from the center of x-y membrane bilayer plane in the Z direction. In Simulation #5, an effector-interacting orientation of K-Ras (where helix 3 and helix 4 are in direct contact with the membrane) was used as the starting structure (Li et al., 2017; Prakash et al., 2016a). In Simulation #1, #2 and #5, the starting configurations of C-Raf^RBD-CRD^ was that of SC0 (see section above). In Simulation #3 and #4, two other different configurations extracted from the 1 μs simulation of C-Raf in solution were adopted. In this way, the initial displacement of CRD relative to the membrane could be larger (~8.5 nm, Simulation #3) or smaller (~5 nm, Simulation #4) than in Simulation #1 and Simulation #2 (~6 nm). In addition, one simulation, #6, was carried out by placing the CRD at the switch II region of K-Ras to test whether this proximity leads to the formation of stable contacts. This simulation was performed for only 240 ns. In all simulations K-Ras4B catalytic-domain was linked to the HVR and anchored to the membrane with a C-terminal farnesyl group pre-inserted into the membrane following previous studies (Li et al., 2017; Prakash et al., 2016a). The parameters for the farnesyl group were generated by the CHARMM generalized force field (CGenFF) (Vanommeslaeghe et al., 2012). The model membrane was composed of 360 POPC (Palmitoyloleoyl-phospatidyl-choline) and 90 POPS (1-Palmitoyl-2-oleoylphosphatidylserine) lipid molecules (20 % POPS). The POPS are equally distributed in each leaflet of the membrane. All model membranes were created by the CHARMM-GUI and equilibrated for 100 ns (Wu et al., 2014).

### Simulation Conditions

The modeled structure of C-Raf^RBD-CRD^: K-Ras4B complex was solvated in a box of TIP3P water. A number of water molecules were replaced randomly by Sodium and Chloride to obtain a neutral charge system with a near-physiological ion concentration of 150 mM. The simulation systems were energy minimized for 2000 steps. Sequentially, a short simulation of 0.5 ns was performed with a harmonic restraint on protein heavy atoms and lipid phosphate groups, followed by another 0.5 ns restraint simulation on only the Cα atoms of the proteins. Production simulations were carried out for 1 μs for Simulation #1 to #5 of the modeled C-Raf^RBD-CRD^: K-Ras4B complex. The initial 50 ns of all simulations were performed using the NAMD/2.10 package (Phillips JC et al., 2005). The NAMD simulations were performed with a time step of 2 fs, and were coupled with a thermostat at 310 K, and a barostat at 1 bar with a semi-isotropic Langevin scheme. A 1.2 nm cut-off (with force-switching between 1.0 and 1.2 nm) was used for van der Waals and for local electrostatic interactions. The long distance electrostatic interaction was treated by the Particle-Mesh Ewald (PME) method. The equilibrated NAMD simulations were converted to Anton2—a specialized supercomputer for molecular dynamics simulation (Shaw et al., 2014), and run on this computer for the remaining 950 ns simulation.

### Free Energy Calculations

In order to calculate the potential of mean force (PMF) of K-Ras4B or C-Raf CRD binding to the membrane, umbrella sampling simulations were performed (Roux et al., 1995). The reaction coordinate is set as the distance in the Z direction between the center of mass of the model membrane and the center of the mass of K-Ras4B helix 4 (residues 127 to 137), or the center of mass of a previously identified lipid binding loop (residues 143 to 151) of the CRD. The membrane is composed of 176 POPC and 44 POPS lipid molecules. The relative orientation of K-Ras4B is set with the helix 4 axis facing the membrane, and CRD is placed with the 143-RKTFLKLAF-151 segment also in parallel facing the membrane. Umbrella windows with a 0.1 nm (Δr) interval were extracted from the simulation trajectories of two steered molecular dynamics (SMD) simulations using established protocols. K-Ras4B was pulled from a position 5.6 nm distant from the membrane center to the membrane surface (Z = 2.2 nm), and then was pulled back in the opposite direction. CRD was pulled from a pre-inserted state (located at Z = 2.0 nm, as the final structure of membrane inserted CRD in simulation #1) to a non-membrane associated position (Z = 5.0 nm), and was pulled back in the opposite direction. A constant pulling velocity of 10^-7^ nm per time step (1fs) was used (total 30-40 ns of each SMD simulation). Biased simulations were performed with a harmonic potential applied between the Cα atoms of selected protein groups (K-Ras4B helix 4 or CRD lipid binding loop) relative to the sliding z-coordinate with a force constant K of 10 kcal/(Mol.Å^2^), typical for such simulations. For K-Ras4B, the sampling simulations were performed for each window for 20 ns at least. For CRD, sampling simulations were applied longer for each window for 60 ns at least, as CRD is partially inserted into the membrane at the lowest energy state. The last 10 ns (K-Ras4B) and 20 ns (CRD) of umbrella sampling trajectory were used in order to calculate the PMF, with the application of the weighted histogram analysis method (WHAM) (Kumar et al., 1992). Uncertainty estimates are determined using the equation 1 of Zhu and Hummer, 2011. The variance in free energy estimators is given by 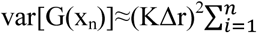 var[x_i_], n is n^th^ simulation window. var[x_i_] is squared error in the estimate of the mean position of x in window i. Variance is estimated using the block averaging method of Flyvbjerg and Petersen, 1985. The standard deviation σ(var[G(x_n_)]) is the cumulative statistical error as shown in Fig. 6b. In total, these simulations amounted to 3.6 μs of trajectories. Although the sampling within each window is longer than used in other recent publications of proteins in and at membranes (e.g. Vunnam et al., 2017) very substantial configurational reorganizations may not be sampled, a standard caveat using a 1D reaction coordinate.

### Analysis

Unless stated otherwise, we performed all our analysis based on the Anton simulations (950 ns) after excluding the initial 50 ns from each trajectory as equilibration. The trajectories of C-Raf^RBD-CRD^ were clustered using Wordom (Fig. 1) (Seeber et al., 2007). The clustering were based on the RMSD using a quality threshold-like algorithm and a cut-off distance of 5 Å. Fusion Protein Modeller (Pham et al., 2007) was used to explore the configurational space of C-Raf^RBD-CRD^ by rotating residues in the shorter linker (132 to 137) (Fig. S1b). The contact maps between one protein domain with another (RBD: CRD, RBD: RAS, CRD: RAS) were made by counting the contact events, i.e. when the distance between residues of one domain and residues of another domain is less than 5.0 Å. Occupancy is the % number of simulation frames where the interaction is present (Fig. S1c and Fig. S3). The criteria for a cation-π interaction is that the distance between all the aromatic ring atoms and the choline nitrogen are below 7 Å and that there is no more than a 1.5 Å in difference between these distances (Fig. S6). For presence of hydrogen bonds a distance of 3.5 Å and angular cut-off of 30 degree was used.

## Acknowledgement

This work is supported by NIGMS grant R01GM112491 to the Buck lab.; P. Prakash is supported by Cancer Prevention and Research Institute of Texas (CPRIT No. DP150093). We thank A.A. Gorfe for helpful discussion and the Ohio Super Computer center located in Columbus, Texas Advanced Computing Center (TACC) and the Extreme Science and Engineering Discovery Environment (XSEDE; Project: MCB150054) for computational resources. Anton2 Computer time was provided by the Pittsburgh Supercomputing Center (PSC) through Grant R01GM116961 from the National Institutes of Health.

## Author contribution

Z.L.L. and M.B. conceived and designed the study with P.P contributing simulation #6; Z.L.L. and P.P. performed the simulations; Z.L.L., P.P. and M.B. analyzed the data; Z.L.L., P.P. and M.B. wrote the manuscript.

